# High-Sensitive Spatial Proteomics for Pancreatic Cancer Progression Analysis

**DOI:** 10.1101/2025.05.01.651678

**Authors:** Jongmin Woo, Zhenyu Sun, Yingwei Hu, Trung Alvin Hoàng, Decapite Christine, Smith Katelyn, Singhi Aatur, Randall E. Brand, Daniel W. Chan, Qing Kay Li, Ralph H. Hruban, Hui Zhang

## Abstract

Pancreatic cancer remains as one of the most challenging malignancies to diagnose and treat due to the late development of symptoms and limited early diagnostic options. Intraductal papillary mucinous neoplasms (IPMNs) are non-invasive precursors to invasive pancreatic ductal adenocarcinoma (PDAC)and an understanding of the changes in patterns of protein expression that accompany the progression from normal ductal (ND) cell, to IPMN to PDAC may provide avenues for improved earlier detection. In this study, we present an optimized spatial tissue proteomics workflow, termed SP-Max (Spatial Proteomics Optimized for Maximum Sensitivity and Reproducibility in Minimal Sample), designed to maximize protein recovery and quantification from limited laser micro dissected (LMD) samples. Our workflow enabled the identification of more than 6,000 proteins and the quantification of over 5,200 protein groups from FFPE tissue contours of pancreatic tissues. Comparative analyses across ND, IPMN, and PDAC revealed critical molecular differences in protein pathways and potential markers of progression. SP-Max provides a systematic, reproducible approach that significantly enhances our ability to study precancerous lesions and cancer progression in pancreatic tissues at unprecedented resolution.

## Introduction

Pancreatic cancer remains one of the most lethal malignancies worldwide, with a five-year survival rate of only 13%^1-5^. The dismal prognosis is largely due to advanced-stage that most tumors are detected, as early symptoms are non-specific or absent and effective screening methods are unavailable. Among the precancerous lesions of the pancreas, intraductal papillary mucinous neoplasms (IPMNs) are relatively common, and are large enough to be detected on imaging)^6-8^. Investigating the molecular landscape of IPMN provides a unique opportunity to elucidate the mechanisms of pancreatic cancer progression and to identify potential biomarkers for early detection and targeted therapies. Traditional proteomic studies in pancreatic cancer have predominantly relied on bulk tissue analysis^9-11^. While these approaches have illuminated key pathways, such as KRAS signaling, WNT/β-catenin signaling, and extracellular matrix remodeling, they lack the spatial resolution required to analyze PDAC and IPMN due to the low neoplastic cellularity of the former. Consequently, the biological complexity underlying tumor microenvironments and the transition from precancerous to invasive states remains underexplored.

Recent advancements in spatial proteomics, particularly those leveraging laser microdissection (LMD)- based workflows, have revolutionized our ability to profile tissue-specific proteomes^12-14^. These techniques enable the isolation of precise tissue regions, allowing for the characterization of protein expression patterns at a localized level. Studies employing LMD-based proteomics have successfully delineated protein signatures associated with tumor progression, glycosylation pathways, and extracellular matrix dynamics in various cancers, including breast and lung^15-19^. Despite their promise, the application of these workflows to pancreatic cancer, particularly in formalin-fixed paraffin-embedded (FFPE) samples, has been limited by challenges such as low cellularity, tissue heterogeneity, and variability in proteomic sensitivity. Emerging methodologies, such as single-cell Deep Visual Proteomics (scDVP), have pushed the boundaries of spatial resolution, achieving single-cell-level insights into tumor microenvironments^20^. However, these approaches typically require fresh or frozen tissue samples to maintain protein integrity, rendering them less accessible for archival FFPE tissues commonly used in clinical research. Moreover, existing FFPE-compatible workflows often struggle with reproducibility and depth of protein identification, underscoring the need for further technical optimization.

This study introduces SP-Max (Spatial Tissue Proteomics Optimized for Maximum Sensitivity and Reproducibility in Minimal Samples), a robust workflow specifically designed to overcome these limitations. By systematically evaluating the effects of tissue section thickness, number of cells, and staining methods, SP-Max achieves unparalleled sensitivity and reproducibility in FFPE tissue analysis. The methodology allows for the detection of over 6,000 proteins and the quantification of more than 5,200 protein groups per tissue section, using minimal sample input. Importantly, SP-Max provides a scalable and reproducible framework for analyzing IPMN, PDAC, and normal duct (ND) regions, facilitating the discovery of region-specific proteomic markers and molecular pathways. This work represents a significant advancement in the field of clinical proteomics for the analysis of PDAC progression, bridging the gap between technical innovation and clinical application. By leveraging SP-Max, we aim to establish a deeper understanding of pancreatic cancer progression, identify early diagnostic markers, and pave the way for the development of personalized therapeutic strategies.

## Methods

### Pancreatic precancerous tissue sample collection

Pancreatic tissue specimens, including normal duct (ND), intraductal papillary mucinous neoplasm (IPMN), and pancreatic ductal adenocarcinoma (PDAC), were acquired from surgically resected samples. Immediately following surgical excision, tissues were fixed in 10% neutral-buffered formalin at room temperature for durations ranging from 12 to 24 hours, depending on tissue size, to ensure thorough preservation of cellular morphology and protein integrity. After fixation, tissues underwent sequential dehydration steps through ascending concentrations of ethanol (70%, 80%, 95%, and 100%, each for one hour), followed by clearing in two changes of xylene (one hour per change) to facilitate complete paraffin infiltration.

Processed tissues were subsequently embedded into molten paraffin wax at approximately 60°C under vacuum conditions in multiple changes (three changes, each for one hour), ensuring complete tissue infiltration and optimal embedding orientation for histological accuracy. Solidified FFPE tissue blocks were then sectioned at a thickness of 5 μm using a rotary microtome. Sections were floated on a heated water bath (40°C) to eliminate wrinkles, collected onto polyethylene naphthalate (PEN)-coated membrane slides (Thermo Fishers), and dried overnight at 37°C to promote optimal tissue adherence suitable for laser microdissection (LMD).

### Deparaffinization and staining FFPE tissue section

FFPE tissue sections on PEN membrane slides were deparaffinized to remove paraffin from the tissue slides. Briefly, FFPE slides were placed in a container with 100% xylene solution for 10 minutes, then transferred to a fresh container with new xylene solution for an additional 10-minute incubation. For the hydration process, deparaffinized slides were sequentially immersed in 100%, 70%, and 50% ethanol, followed by water, for five minutes each. Before staining, the slides were equilibrated with 1x PBS buffer.

For Hematoxylin staining, the PBS buffer on the slide was gently removed by tapping the slide on a paper towel. Then, 2 mL of Hematoxylin (Cat# MHS1-100ML, Sigma Aldrich) was applied to cover the tissue, and the slide was incubated for 3 minutes. After rinsing with distilled water, the stained tissue samples were briefly dipped in 0.5% Ammonium Hydroxide for 1 minute to achieve bluing, followed by a final rinse with distilled water. For Toluidine Blue O (TBO) staining, deparaffinized pancreas tissue sections were covered with a dropwise application of 0.5% w/v TBO (Cat# 1159300025, Sigma Aldrich) for 1.5 minutes. Excess TBO stain was removed, and the sections were sequentially rinsed with distilled water for 1 minute. Unstained tissue sections were prepared by gently removing excess PBS by washing with distilled water. All samples were dried by high-purity nitrogen before utilizing.

### Estimation of Equivalent Cell Counts in Tissue Sections

To estimate cellular equivalents within each captured area, the volume of individual epithelial cells (∼4,000 μm^3^) was used as a reference^21, 22^. Tissue volume was calculated by multiplying capture area by section thickness (e.g., 5 μm, 10 μm, or 20 μm). For example, a 40 μm × 40 μm area from a 5 μm-thick section corresponds to the equivalent of two cells. Larger areas, such as 420 μm × 420 μm, correspond to approximately 200 cells. This calculation standardizes sample input for reproducible proteomic analysis across experiments.

### Laser microdissection

The stained or unstained tissue sections on PEN membrane slides were thoroughly dried at room temperature before proceeding with LMD. The slides were placed on the slide rack of an LMD-7000 (Leica, US), ROIs in the FFPE tissue slide were identified at 20x magnification. The laser control parameters were set as follows at 20x magnification: power of 54, aperture of 1, speed of 5, specimen balance of 50, head current of 100%, pulse frequency of 773, and offset of 110, to prepare for precise tissue voxel dissection. The ROI tissue sections were collected in the flat cap of a 0.2 mL PCR tube (Cat# 6571, Corning, Tamp, MEX) containing 20 μL of DMSO. To ensure accurate collection of small tissue sections, the cap was carefully inspected after each LMD operation. The tubes were then centrifuged at 15,000 xg for 1 minutes, and the samples were dried in a speed vacuum for 4 hours, ensuring complete removal of any remaining liquid.

### Proteomics sample preparation

We added 1.5 μL of a lysis buffer mixture, containing 0.1% n-Dodecyl-β-D-Maltoside (DDM), 0.02% Lauryl Maltose Neopentyl Glycol (LMNG), 5 mM TCEP, and 0.001unit DNase in 100 mM Triethylammonium Bicarbonate (TEAB), to the sample and incubated it at 80°C for 1 hour, with a heating lid set to 100°C. After a 5-minute sonication, the samples were cooled to room temperature and were sequentially digested by adding 0.5 μL of 50 ng/μL LysC (Waco, US) and an equal amount of 50 ng/μL Trypsin (Promega, US), allowing the enzymes to digest the sample overnight at 37°C, with a heating lid set to 57°C. The samples were then acidified by adding 0.5 μL of 5% formic acid and incubated for 10 minutes at room temperature. Finally, peptides were transferred and loaded onto the pre-conditioned Evotips (Evosep, US) following the’s instructions.

### Proteomic analysis using LC-MS instrument

Proteomic analyses were conducted using DIA-MS on two distinct instruments paired with the Evosep One LC: Orbitrap Ascend (Thermo) and timsTOF HT (Bruker). Peptide samples prepared from tissue sections of varying staining, size, and thicknesses were separated on the Evosep using an 88-minute gradient, allowing for the analysis of up to 15 samples per day (SPD) with an analytical column (Cat# 1893474, PepSep C18, 15 cm x 150 μm, 1.5 μm). For the Orbitrap Ascend, which utilizes Field Asymmetric Ion Mobility Spectrometry (FAIMS), the analysis settings included a compensation voltage of −55 V, a carrier gas flow rate of 3.5 L/min, a mass range of 380-1,000 m/z, a variable isolation window with 17 scan events, an MS2 scan range of 145-1,450 m/z, a resolution of 30,000, an Automatic Gain Control (AGC) target of 1×10^6, and a maximum injection time of 200 ms. Peptide samples from individual dissected pancreas tissue sections representing different disease conditions were separated using the Whisper Zoom 40 SPD method (33 minutes gradient) on the Evosep One with an analytical column (Cat# 1893473, PepSep C18, 15 cm x 75 μm, 1.9 μm). These samples were then analyzed on the timsTOF HT with settings as follows: a mobility range of 0.70 – 1.45 1/k0, a mass range of 100 – 1,700 m/z, and an accumulation time of 166 ms with high-sensitivity detection enabled. Additionally, the collision energy was set to 20 eV at 0.60 V·s/cm^2^ and 59 eV at 1.60 V·s/cm^2^. For DIA-PASEF, mass width was set as 25 Da ranged from 338.6 Da to 1338.6 Da, resulting in total 40 mass step per cycle and 1.72 s estimated cycle time.

### Data and statistical analysis

Proteomic raw data for SP-Max performance assessment in varying conditions or for pancreatic cancer study using FAIMS-Ascend or timsTOF HT, respectively, were processed using Spectronaut (version 19.0.24) with the direct-DIA approach (FASTA database downloaded 09/15/2023, UniprotKB reviewed, 20,426 entries). Briefly, raw DIA data were directly used for analysis using non-cross-normalized mode. The following processing parameters were used for all software tools: acetylation at the protein N-terminus (Acetyl [N-term]) and oxidation of methionine residues (Oxidation [M]) were included as variable modifications with a maximum of two missed cleavages, while cysteine carbamidomethylation (+57.0215 Da) as the fixed modification. FDR was set to 1% using the target-decoy strategy on both the peptide and protein levels.

Statistical significance was assessed using rigorous methods to ensure the reliability of the results using Perseus (version 1.6.15). Quantitative values were log2-transformed, and protein groups were identified if quantitative values were present. For protein quantification, proteins were filtered by quantified proteins if quantified at least 70% of the samples in total experimental group contained valid measurements. No normalization was performed for the data of the SP-Max method assessment in different conditions, while the width adjustment normalization was conducted for the pancreatic cancer dataset. Missing values were then imputed from normal distribution with a width of 0.3 and a down shift of 1.8. Batch effects from individual sample or different localized area on the tissue section were removed by ComBat batch effect correction algorithm. A multiple sample ANOVA among three groups (ND, IPMN, and PDAC) with a significance threshold of FDR < 0.01 and S^0^=0.1was applied to identify differentially expressed proteins.

### Bioinformatic analysis

For Gene Ontology (GO) enrichment analysis, significantly regulated proteins (ANOVA FDR < 0.01, fold change > 2) were uploaded to MetaScape as input, annotated as Homo sapiens genes^23^. Enrichment for GO biological processes was performed, and results with a p-value < 0.05 were considered statistically significant. Single-sample Gene Set Enrichment Analysis (ssGSEA) was performed using GenePattern to evaluate Reactome pathway activity^24, 25^. The MSigDB dataset (C2.Reactome.v2024.1.HS.symboles.gmt) was used as a reference for pathway annotation. Gene sets with an overlap of more than 10 genes from the analysis were considered for further interpretation, focusing on pathways relevant to pancreatic cancer progression. In order to verify the pancreatic cell population within the LMD-contoured samples, we compared the identified proteins against the single-cell transcriptome atlas of the human pancreas^26^. This atlas provides a comprehensive classification of pancreatic cell types, detailing five distinct cell types within the endocrine compartment (alpha, beta, delta, epsilon, and pancreatic polypeptide [PP] cells) and four within the exocrine compartment (ductal, acinar, mesenchymal, and endothelial cells). This comparison enabled precise validation of the cellular composition in our samples, ensuring accurate representation of ductal epithelial cells while minimizing potential contamination from other pancreatic cell types.

## Results

### Streamline SP-Max workflow for high-sensitive spatial tissue proteomics

To enhance the sensitivity and reproducibility of spatial proteomics in FFPE pancreas tissues, we developed the SP-Max workflow (Figure 1). This optimized workflow integrates multiple innovations to address the challenges of analyzing limited and heterogeneous clinical samples, providing a systematic method to achieve high-quality proteomic data from FFPE tissues. The process begins with Hematoxylin and Eosin (H&E) staining, which enables precise annotation of regions of interest (ROIs) by expert pathologists using high-resolution imaging. These ROIs, carefully selected to distinguish normal duct (ND) and disease stages, are subsequently dissected from adjacent tissue sections using laser microdissection (LMD). The staining approach was specifically optimized to ensure compatibility with proteomics workflows, balancing efficient visualization of tissue architecture with minimal interference in downstream LC-MS analysis. Thorough evaluations of slide thickness and sample size were performed, leading to an optimized protocol that ensures high reproducibility and sensitivity even for minimal sample input. This systematic optimization allows for robust spatial proteomic profiling of pancreatic tissues, overcoming the inherent challenges of FFPE samples.

**Figure 1.**
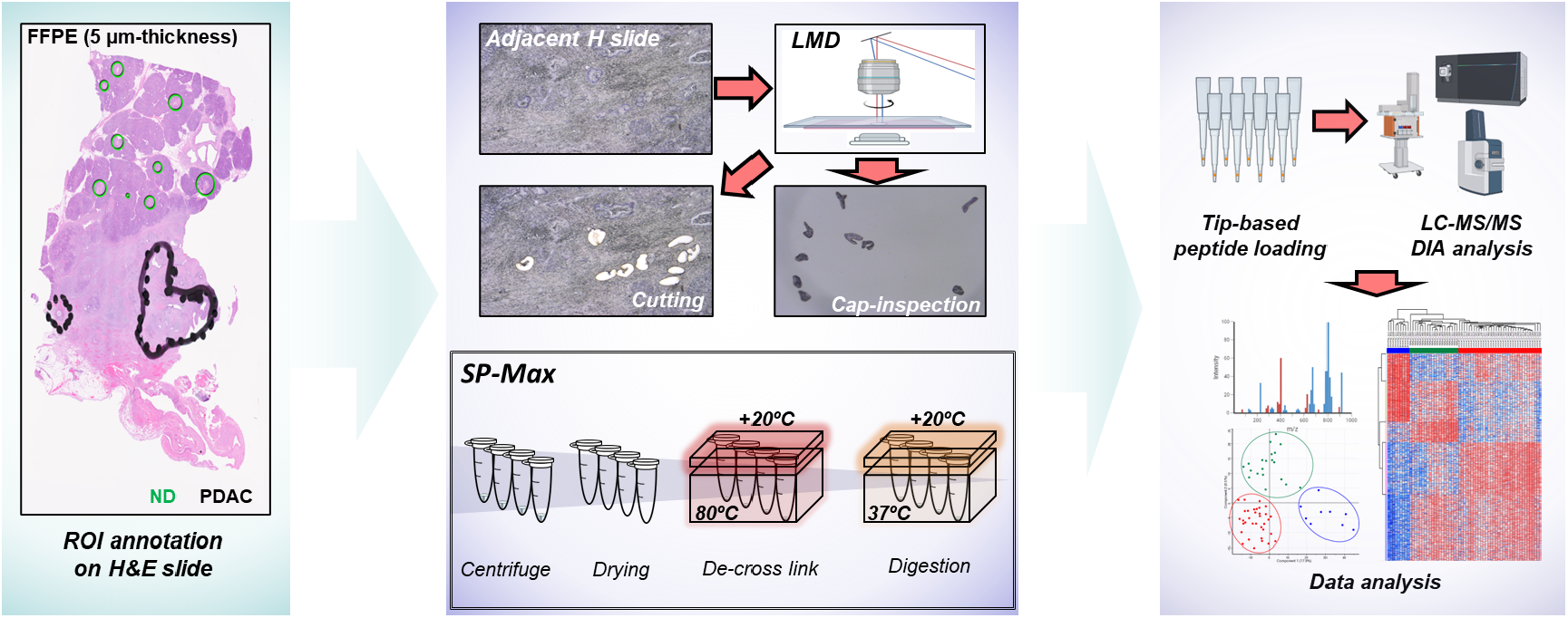
Workflow of LMD-based FFPE tissue preparation for high-sensitivity spatial proteomics. Regions of interest (ROIs) were annotated on hematoxylin and eosin (H&E)-stained FFPE tissue sections to differentiate between normal duct (ND) and distinct disease stages. Laser microdissection (LMD) was utilized to precisely isolate these ROIs, with careful slide and cap inspection to ensure proper sample collection from LMD. The SP-Max workflow includes optimized steps to minimize sample loss, particularly through the application of a temperature gradient (+20°C) during the de-cross linking and enzymatic digestion process, enhancing protein recovery. Dissected tissues underwent sonication for tissue lysis and were subjected to controlled heating (80°C for de-cross linking followed by 37°C for digestion). Resulting peptides were loaded on Evotips and analyzed via LC-MS/MS using a data-independent acquisition (DIA) approach. Subsequent data analysis provided high-resolution proteomic patterns across tissue regions, enabling comprehensive spatial proteomics.

The SP-Max workflow was further refined to maximize the efficiency of sample preparation while minimizing sample loss, especially critical for limited sample quantities. For example, during the de-crosslinking step, temperature gradients were strategically applied, with the vessel lid maintained at 100°C and the sample itself heated to 80°C. This method effectively prevents evaporation and ensures consistent sample volume retention, which is critical when working with a few microliters of material. By preserving sample integrity during lysis and digestion, this approach significantly enhances peptide yield and digestion efficiency. These adaptations reduce the labor-intensive nature of traditional methods, enabling streamlined processing of limited clinical samples with minimal hands-on effort.

The peptides generated through SP-Max are processed via C18-based Evotip loading to minimize handling loss and analyzed using data-independent acquisition mass spectrometry (DIA-MS). This comprehensive workflow delivers robust and reproducible data, providing detailed molecular profiles of ND, IPMN, and PDAC tissue regions. By integrating these optimizations, SP-Max establishes a reproducible, high-sensitivity platform for FFPE-based spatial proteomics, facilitating the discovery of critical molecular pathways and potential biomarkers for pancreatic cancer progression.

### Evaluation of tissue staining, areas, and thickness for reliable spatial proteomic analysis

In clinical research, especially studies involving pancreatic cancer, the limited cellularity of the neoplastic cells and of the non-neoplastic ductal cells in tissue samples require careful optimization of sample use to ensure reliable data. Additionally, tissue heterogeneity influenced by section thickness can affect the specificity of proteomic data, making it essential to evaluate the optimal sample size and thickness for reproducible protein detection. To further optimize the SP-Max workflow, we evaluated different staining methods to identify the most suitable approach for proteomic analysis (Figure 2A, Supplementary Table S1). Our results showed that Hematoxylin staining provided the best performance in terms of protein identification compared to Toluidine Blue O (TBO) and unstained samples (Figure 2B). Consistent with our previous study^13^, Hematoxylin, a non-binding reagent to proteins, effectively visualized ROIs without interfering with downstream proteomic analysis, making it the preferred choice for further evaluations involving sample size and thickness.

**Figure 2.**
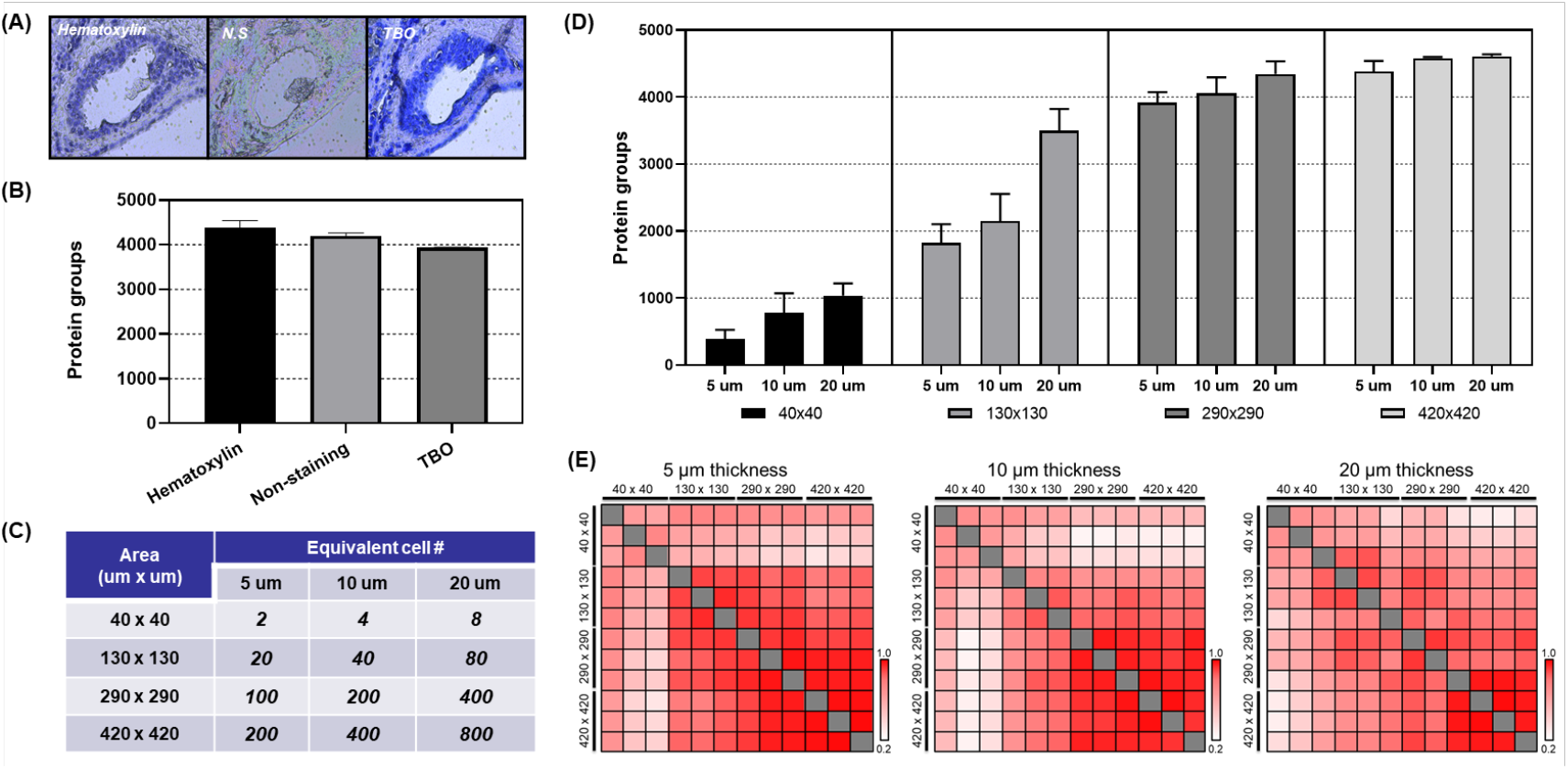
Tissue Staining, areas and thickness for reliable spatial proteomic analysis (A) Tissue slides stained with Hematoxylin, Toluidine Blue O (TBO), or left unstained (NS). (B) Number of protein groups identified from a 420 μm × 420 μm tissue area with 5 μm thickness using different staining or non-staining methods. (C) Estimated equivalent cell counts based on sample volume (μm^3^) for various tissue areas and thicknesses, assuming a single cell is 20 μm diameter and volume of approximately 4,000 μm^3^. (D) Number of identified protein groups across different tissue sample sizes and amounts, varying in area and thickness. (E) Heatmap illustrating correlation between biological replicates and between scaled sample sizes, measured across three different tissue thicknesses (5 μm, 10 μm, and 20 μm).

Next, we assessed tissue sections of varying thicknesses (5 μm, 10 μm, and 20 μm) and sample sizes ranging from two cells to approximately 800 cells captured via LMD (Figure 2C, Figure S1A and S1B, Supplementary Table S2). This assessment was crucial for understanding how these variables influenced protein recovery and detection fidelity, particularly when using the SP-Max workflow in combination with archival FFPE samples. Our data indicated that increased sample input correlated with greater proteome depth (Figure 2D). Specifically, when the sample input increased from 40 cells (130 μm-side length square area in 10 μm-thickness) to 80 cells (130 μm-side length square area in 20 μm-thickness), the number of identified proteins rose from 2,147 to 3,496, respectively, showcasing an 163% increase in protein identification efficiency. This increasement can be attributed to enhanced tryptic digestion efficiency, as the reaction involved 50 ng of enzymes in 2.5 μL of buffer on the SP-Max, suggesting that smaller initial sample amounts may have limited digestion efficiency.

Additionally, we observed heterogeneity within cell populations in samples of the same size by correlation of quantified proteins among triplicate biological samples (Figure 2E). Thicker tissue sections required lager area size to achieve high reproducibility (Pearson correlation r ≈ 0.98) due to the increased heterogeneity of tissue in three dimensions, which could introduce variability. In contrast, thinner sections (e.g., 5 μm) provided clearer proteomic signals with smaller area size, reducing potential contamination from neighboring areas and resulting in higher data consistency. The optimal conditions for reproducible results were found to vary based on tissue thickness. Thinner sections (e.g., 5 μm) needed smaller area (as few as 20 cells) to achieve high reproducibility, whereas thicker sections (e.g., 20 μm) required larger area size (800+ cells) to maintain consistency due to greater sample heterogeneity. These observations highlight the trade-off between tissue thickness and sample size: while thinner sections yield more reproducible data with fewer cells, thicker sections, although richer in protein content, demand larger sample sizes to counteract increased heterogeneity and variability. The findings from our evaluations underscore that SP-Max is highly efficient for detecting a broad range of proteins, particularly as sample input increases. However, the reproducibility of proteomic analysis can be influenced by the sample’s thickness and size, necessitating tailored approaches depending on the available tissue and study objectives.

### Characterization of pancreatic cancer and IPMN proteome using the optimized spatial proteomics workflow

To characterize the proteomic landscape of pancreatic cancer progression, we applied the SP-Max workflow for clinical FFPE samples to analyze protein expression profiles across normal duct (ND), intraductal papillary mucinous neoplasm (IPMN), and pancreatic ductal adenocarcinoma (PDAC) samples collected via LMD. A total of 10 cases were analyzed, with 17 marked tissue features corresponding to specific disease conditions. More than three biological replicates were captured from each tissue feature, resulting in the identification of over 6,900 proteins and the quantification of over 5,200 proteins (Supplementary Table S3). These proteins were derived from a 290 μm-side length square area at 5 μm thickness, equivalent to approximately 100 cells (Figure 3A). To ensure sample purity, the contours of pure ductular epithelial cells were meticulously delineated and collected with guidance from a pathologist. However, minor contamination from adjacent cell types, such as acinar or stromal cells, may have influenced the proteomic profiles. To evaluate cellular purity, we employed cell-type-specific signature proteins identified from a single-cell transcriptome atlas of the human pancreas. These signature proteins encompassed markers for diverse pancreatic cell populations, including ductal epithelial, acinar, endothelial, alpha, beta, delta, epsilon, polypeptide (PP), and mesenchymal cells (Figure 3B). Analysis confirmed that the majority of the cell population in our LMD dataset corresponded to ductular epithelial cells, with acinar cells constituting the next most prominent group. These findings highlight the high quality of the tissue contours obtained through LMD. Additionally, principal component analysis (PCA) revealed distinct clustering of ND, IPMN, and PDAC samples. PDAC and IPMN formed closely related clusters, reflecting shared molecular features, while ND samples formed a separate and distinct cluster (Figure 3C). A total of 1,649 significantly regulated proteins were identified across the three groups through ANOVA (FDR < 0.01, S0 = 0.1) and categorized into five distinct expression clusters (Figure 3D, Supplementary Table S4). Cluster analysis revealed critical molecular transitions, including the upregulation of proteins implicated in tumorigenesis and cancer progression in IPMN and PDAC compared to ND.

**Figure 3.**
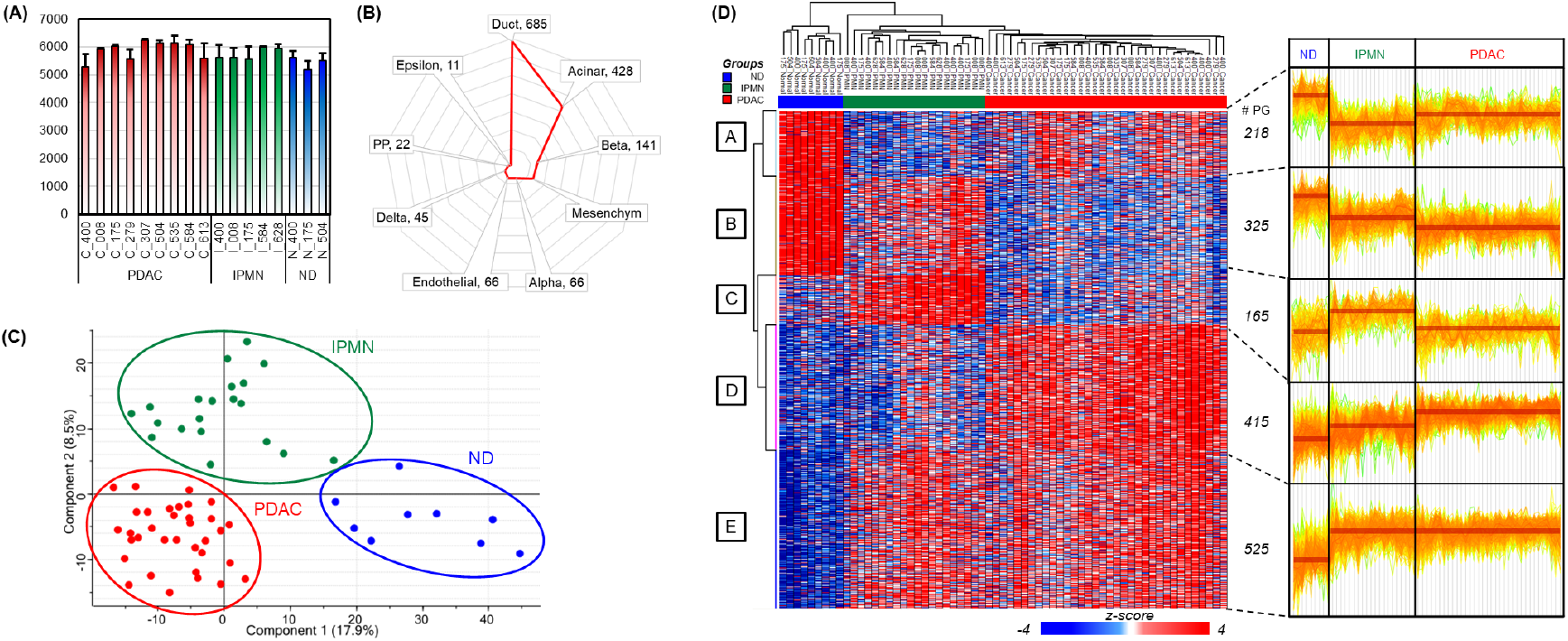
Characterization of pancreatic cancer and IPMN proteome. (A) Protein group counts identified across individual samples from PDAC (red), IPMN (green), and ND (blue) tissue sections. A total of 10 cases were analyzed, with 17 marked tissue features corresponding to specific disease conditions. At least three biological replicates were captured from each feature. (B) Cell-type-specific signature proteins derived from a single-cell transcriptome atlas of the human pancreas^26^, highlighting key markers in ductal, acinar, beta, mesenchymal, and other cell types. (C) Principal component analysis (PCA) showing distinct clustering of ND, IPMN, and PDAC tissue samples. PDAC and IPMN, representing cancerous and precancerous tissues of pancreas, respectively, cluster closely, while ND forms a separate group. (D) Heatmap of 1,649 significantly regulated proteins identified by ANOVA (FDR < 0.01, S0 = 0.1), clustered into five distinct expression patterns (labeled A–E). The right panel shows expression profiles of each cluster across ND, IPMN, and PDAC LMD-tissue lesions.

### Biological processes Enriched in regulated proteins in IPMN and PDAC

To elucidate the molecular mechanisms underlying IPMN and PDAC progression, biological processes enriched in upregulated (Cluster E in Figure 3D) and downregulated (Cluster B in Figure 3D) proteins were analyzed (Figure 4A-B, Supplementary Table S5). Proteins in Cluster E, upregulated in both IPMN and PDAC compared to ND, were associated with several critical biological processes (Figure 4A). Ribosome biogenesis emerged as the most significantly enriched process, underscoring the enhanced protein synthesis capacity that is critical for sustaining the rapid proliferation of cancer cells. This finding aligns with previous studies reporting that ribosomal protein overexpression is linked to tumorigenesis and poor prognosis in various cancers^27-31^. Similarly, the enrichment of chromosome condensation highlights the elevated mitotic activity characteristic of cancer progression. The observed enrichment in heterophilic cell-cell adhesion points to alterations in cell adhesion dynamics, which may facilitate tumor cell invasion and metastatic dissemination—a hallmark of advanced cancers. Iron ion transport and mitochondrial respirasome assembly were also significantly enriched, reflecting the metabolic reprogramming and elevated bioenergetic demands of cancer cells. These biological processes are well-documented contributors to the metabolic flexibility of tumors, enabling them to thrive under hypoxic and nutrient-limited conditions^32-34^. Furthermore, the negative regulation of apoptotic signaling pathways and regulation of cellular response to stress highlight the ability of IPMN and PDAC cells to evade cell death and adapt to the hostile tumor microenvironment. The enrichment of insulin response processes suggests potential crosstalk between metabolic and proliferative pathways, consistent with the role of insulin signaling in promoting cancer cell survival and growth. Together, these processes underscore the multifaceted molecular adaptations that underlie the aggressive phenotypes of IPMN and PDAC.

**Figure 4.**
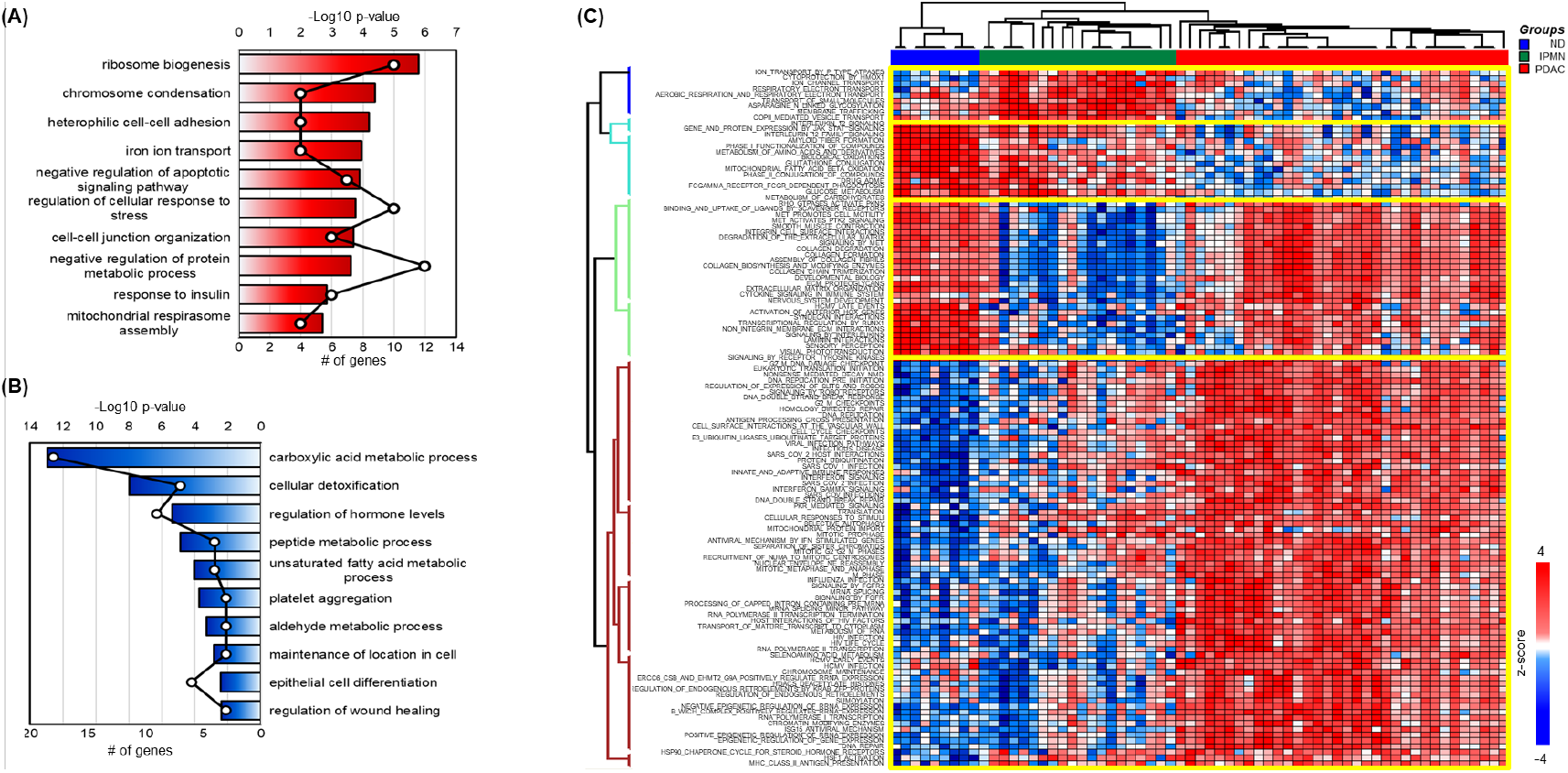
Biological process enrichment and pathway analysis of protein clusters in pancreatic cancer progression. (A) Biological processes (BPs) enriched in proteins from Cluster E in Figure 3D, which are upregulated in both IPMN and PDAC. A total of 138 proteins with a ≥2-fold change compared to normal duct (ND) were analyzed. (B) Biological processes enriched in proteins from Cluster B in Figure 3D, which are downregulated in IPMN and PDAC. A total of 70 proteins with a ≥2-fold change compared to ND were analyzed. (C) Single-sample gene set enrichment analysis (ssGSEA) of all proteins from Figure 3D, demonstrating the enrichment of Reactome pathways for individual tissue samples. The enrichment scores for each sample are provided in Supplementary Table S6.

In contrast, proteins in Cluster B, progressively downregulated in IPMN and PDAC compared to ND, were enriched in biological processes essential for maintaining normal cellular and tissue homeostasis (Figure 4B). Carboxylic acid metabolic processes and cellular detoxification were the most significantly suppressed processes. The downregulation of these processes aligns with previous findings that metabolic impairments and oxidative stress accumulation contribute to cancer progression^35-38^. Regulation of hormone levels and peptide metabolic processes were also significantly disrupted, reflecting altered intercellular communication and proteostasis, both of which are critical for maintaining normal tissue function. The suppression of processes such as platelet aggregation and wound healing suggests a shift in the tumor microenvironment that favors invasion and metastasis over tissue repair and homeostasis. Additionally, the downregulation of epithelial cell differentiation and maintenance of cellular location highlights the loss of epithelial integrity and polarity, hallmark features of the epithelial-to-mesenchymal transition (EMT). EMT is a well-established mechanism by which cancer cells acquire invasive and metastatic potential, further emphasizing the relevance of these findings in the context of IPMN and PDAC progression^39-42^.

To complement the GO analysis, we conducted single-sample Gene Set Enrichment Analysis (ssGSEA) to quantify pathway-level phenotypes and provide a robust scoring system for each sample, enabling a more refined analysis of Reactome pathways (Figure 4C, Supplementary table S6)^25^. This approach highlights independent pathway dynamics across ND, IPMN, and PDAC, offering quantitative insights into disease progression. Pathways such as aerobic respiration and membrane trafficking were prominently activated in IPMN but became inactivated in PDAC. This pattern, which includes pathways like respiratory electron transport and asparagine N-linked glycosylation, reflects heightened metabolic activity and cellular maintenance in IPMN. Their subsequent suppression in PDAC underscores the metabolic rewiring characteristic of malignant transformation, with a shift towards glycolysis (Warburg effect) and reduced oxidative phosphorylation. This transition highlights metabolic vulnerabilities in PDAC that may serve as therapeutic targets.

In contrast, pathways involved in immune regulation, detoxification, and energy production, including interleukin-12 signaling, glutathione conjugation, mitochondrial fatty acid beta-oxidation, and carbohydrate metabolism, exhibited progressive inactivation from ND to IPMN and PDAC. These findings emphasize a progressively immunosuppressive and metabolically compromised tumor microenvironment as the disease advances. The diminished interleukin-12 signaling reflects weakened immune surveillance, while reduced glutathione conjugation indicates impaired oxidative stress management, further exacerbating tumor progression. Interestingly, extracellular matrix (ECM)-related pathways such as collagen degradation, integrin interactions, and ECM proteoglycan organization were inactivated in IPMN but reactivated in PDAC. This reactivation highlights the critical role of ECM remodeling in enabling PDAC invasion and metastasis. Pathways like MET activation and Rho GTPase signaling, which promote motility and cytoskeletal reorganization, further underscore the aggressive phenotype of PDAC and the pivotal role of ECM as a therapeutic target. Additionally, pathways related to cell cycle regulation, DNA repair, and RNA processing, such as G2/M DNA damage checkpoints, homology-directed repair, and nonsense-mediated decay, demonstrated gradual activation from ND to IPMN and PDAC. These pathways reflect the genomic instability and proliferative demands of cancer cells, offering potential avenues for therapeutic intervention. The activation of immune-related pathways, such as interferon signaling, likely represents an adaptive response to oncogenic stress, though its precise role in pancreatic cancer progression warrants further investigation.

This study provides novel insights into pancreatic cancer progression by integrating spatial proteomics and ssGSEA pathway analysis. Key findings include the identification of unique pathway transitions, such as the suppression of immune and metabolic pathways in IPMN and their subsequent reactivation in PDAC. The delineation of progressively activated DNA repair and cell cycle pathways highlights their therapeutic relevance, particularly in targeting proliferative and repair mechanisms in PDAC. The reactivation of ECM remodeling pathways underscores their importance in metastatic progression, presenting opportunities for disrupting tumor invasion and dissemination. Together, these findings advance our understanding of pancreatic cancer progression and open new avenues for targeted therapeutic strategies.

### Significantly regulated proteins in IPMNs

To investigate the pathways enriched in precancerous lesions, we focused on the 165 proteins in Cluster C (Figure 3D), which were upregulated in IPMN compared to ND but downregulated in PDAC. This protein set represents highly regulated proteins specific to IPMNs. Reactome pathway analysis of these 165 proteins revealed several significantly enriched pathways, as shown in Figure 5A. Among these, the most significant pathway was the O-linked glycosylation of mucins, a well-known and critical pathway in IPMN, as reflected in the disease’s nomenclature^43^. Our study underscores the importance of this pathway within the context of LMD-based proteomic analysis. The O-linked glycosylation of mucins pathway was enriched with nine key proteins, including B2GNT7, C1GALT1, GALNT1, GALNT4, GALNT5, GALNT6, GALNT7, MUC13, and MUC5AC, each demonstrating significant upregulation in IPMN compared to ND (Figure 5B). Quantification confirmed that these proteins were markedly increased in IPMN, highlighting their potential roles in the formation and progression of mucin-producing neoplasms.

**Figure 5.**
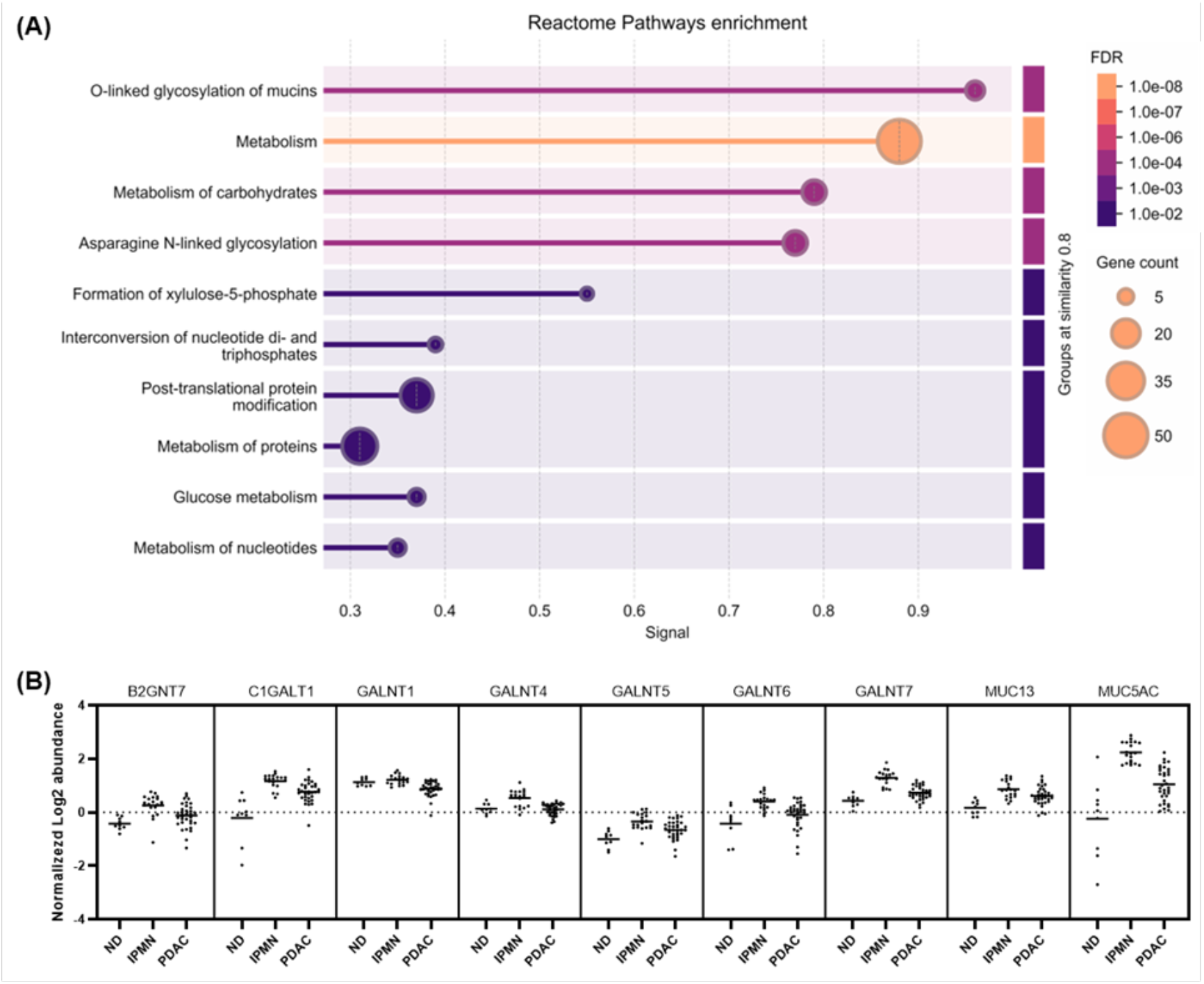
Altered proteins in IPMNs. (A) Bubble plot depicting Reactome pathway enrichment of Cluster C proteins. The size of each bubble represents the number of proteins contributing to the pathway, while the color indicates statistical significance (FDR). (B) Quantitative differences in expression levels between ND, IPMN, and PDAC for the nine proteins enriched in the O-linked glycosylation of mucins pathway. Each protein demonstrates elevated level of proteins in IPMN compared to ND, followed by downregulation in PDAC.

To gain deeper insights into the molecular pathways enriched in precancerous lesions, we focused on proteins upregulated in IPMN compared to ND and PDAC (Cluster C in Figure 3D, Supplementary Table S4), representing key processes in the transition from a precancerous to a malignant state. Several clusters of pathways were identified using ssGSEA, providing biological and clinical insights into the molecular mechanisms of IPMN (Supplementary Figure 2). Pathways related to membrane trafficking and vesicle transport, including Golgi transport, Rab-mediated trafficking, COPII-mediated vesicle transport, and intra-Golgi traffic, were also significantly activated in IPMN. These pathways facilitate the efficient secretion of mucins and other macromolecules, supporting the high secretory activity characteristic of IPMN. This upregulation highlights the importance of maintaining robust cellular machinery to sustain mucin production. However, in PDAC, these pathways were downregulated, reflecting a metabolic shift as cancer cells prioritize processes supporting invasion and metastasis over secretion. Metabolic and energy pathways, such as glucose metabolism, lipid metabolism, respiratory electron transport, and glycerophospholipid biosynthesis, were actively upregulated in IPMN. These pathways provide the energy and biosynthetic precursors required for mucin production and cellular proliferation. The increased metabolic demand in IPMN reflects its unique cellular state as a precancerous lesion with high secretory output. Conversely, in PDAC, the metabolic reprogramming shifts toward pathways that enhance tumor survival under hypoxic and nutrient-deprived conditions, consistent with a malignant phenotype.

## Discussion

The SP-Max workflow represents a significant advancement in the application of spatial proteomics to clinical FFPE tissue samples, addressing longstanding challenges in sample preparation and data reproducibility. By enabling the identification and quantification of over 6,000 proteins from limited input samples, SP-Max demonstrates unparalleled sensitivity, making it a robust method for studying heterogeneous and cellularly sparse tissues such as pancreatic cancers. One of the key strengths of SP-Max is its ability to handle minimal sample quantities without compromising data quality. The systematic optimization of variables such as staining methods, slide thickness, and sample area ensured reproducibility and depth of proteomic coverage. For instance, the use of Hematoxylin staining coupled with precise LMD facilitated the isolation of specific tissue regions while maintaining compatibility with proteomics workflows. Moreover, innovations such as temperature gradient-controlled de-crosslinking and enzymatic digestion minimized sample loss by preventing evaporation, further enhancing peptide yield and digestion efficiency. These features make SP-Max particularly well-suited for the analysis of archival FFPE tissues, which are often the only available samples in clinical research. Recently, wcSOP-MS has been demonstrated to enable reliable label-free quantification in the same size of LMD-based tissue sections^22^. While wcSOP-MS has been applied to FFPE-derived tissue voxels, its reported protein identifications remain lower than those achieved by SP-Max, with approximately 3,000 protein groups detected. This difference may be attributed to the optimized reaction volume used in SP-Max (2.5 μL instead of 10 μL) and the implementation of a heating lid set to an additional 20°C, which collectively enhance digestion efficiency and minimize sample loss. These refinements highlight the importance of precise reaction conditions in maximizing proteomic depth from limited clinical samples.

Despite its strengths, SP-Max also has limitations that should be addressed in future studies. The workflow’s reliance on specialized equipment, such as high-resolution LMD systems and DIA-compatible mass spectrometers, may limit its accessibility in lower-resource settings. Additionally, while the method achieved remarkable sensitivity and reproducibility, the processing of extremely small sample volumes still requires careful handling to avoid variability. Scaling the workflow for high-throughput applications will require further automation and standardization to reduce labor intensity and ensure consistency across multiple operators. To overcome these challenges, future efforts will focus on optimizing sample preparation protocols to enhance robustness and reproducibility, even with minimal input volumes. Additionally, we aim to integrate automated liquid handling systems and microfluidic platforms to streamline sample processing and improve throughput while minimizing human-induced variability^13, 44^. Expanding compatibility with a broader range of mass spectrometry platforms, including emerging cost-effective DIA-compatible systems, will also be prioritized to increase accessibility. These advancements will ensure that SP-Max can be more widely adopted and effectively utilized in diverse research and clinical settings.

The application of SP-Max to pancreatic cancer tissues revealed novel insights into the molecular mechanisms underlying the progression from normal duct (ND) to IPMN and ultimately to PDAC. Notably, the analysis identified significant pathway transitions, including the upregulation of ribosome biogenesis and metabolic reprogramming in IPMN and PDAC, as well as the suppression of immune and detoxification pathways. These findings highlight potential vulnerabilities in pancreatic cancer cells, such as their reliance on oxidative phosphorylation and ribosomal activity, which could be targeted therapeutically. Furthermore, the reactivation of extracellular matrix (ECM) remodeling pathways in PDAC underscores the importance of the tumor microenvironment in facilitating invasion and metastasis^45, 46^. This study also underscores the value of integrating spatial proteomics with pathway-level analyses, such as ssGSEA, to capture the dynamic changes in molecular phenotypes across disease stages. By providing quantitative pathway scoring and single-sample resolution, SP-Max enabled the identification of region-specific proteomic signatures and potential biomarkers for early detection and targeted therapy. These advancements not only enhance our understanding of pancreatic cancer progression but also pave the way for the broader application of spatial proteomics in other challenging cancer types.

In conclusion, SP-Max provides a reproducible and high-sensitivity platform for spatial proteomics, bridging the gap between technical innovation and clinical application. By addressing the limitations of traditional workflows and uncovering novel molecular insights, SP-Max has the potential to transform how precancerous lesions and cancer progression are studied, offering new opportunities for early intervention and personalized treatment strategies.

## Abbreviations

IPMN: Intraductal Papillary Mucinous Neoplasm
PDAC: Pancreatic Ductal Adenocarcinoma
ND: Normal Duct
SP-Max: Spatial Proteomics Optimized for Maximum Sensitivity and Reproducibility in Minimal Sample
LMD: Laser Microdissection
FFPE: Formalin-Fixed Paraffin-Embedded
DIA-MS: Data-Independent Acquisition Mass Spectrometry
FAIMS: Field Asymmetric Ion Mobility Spectrometry
GO: Gene Ontology
ssGSEA: Single-Sample Gene Set Enrichment Analysis
PCA: Principal Component Analysis
TEAB: Triethylammonium Bicarbonate
TCEP: Tris(2-carboxyethyl) phosphine
TBO: Toluidine Blue O
MSigDB: Molecular Signatures Database

## Supplementary Information

**Figure S1.**
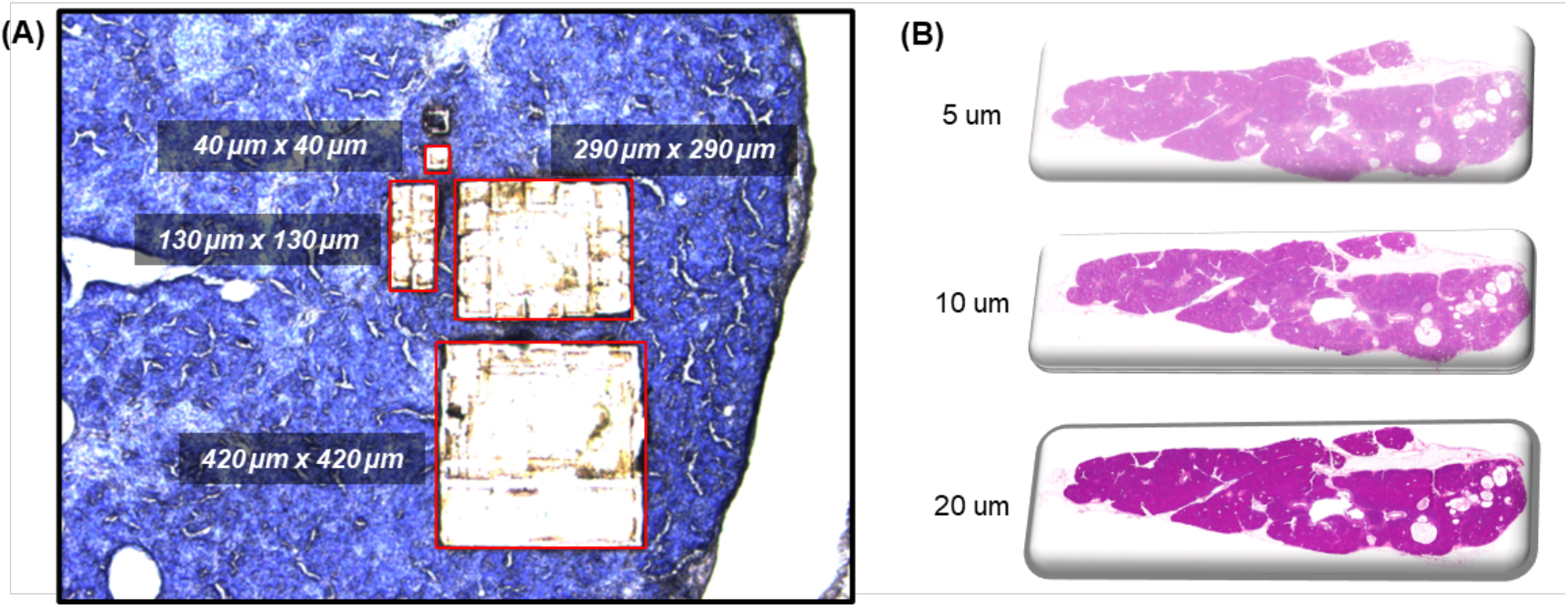
(A) Laser microdissection (LMD) capture areas shown for varying cell counts on a 5 μm thick pancreas tissue section. (B) H&E-stained images of pancreas tissue sections prepared at three different thicknesses: 5 μm, 10 μm, and 20 μm, demonstrating the structural consistency in tissue morphology with slide thickness.

**Figure S2.**
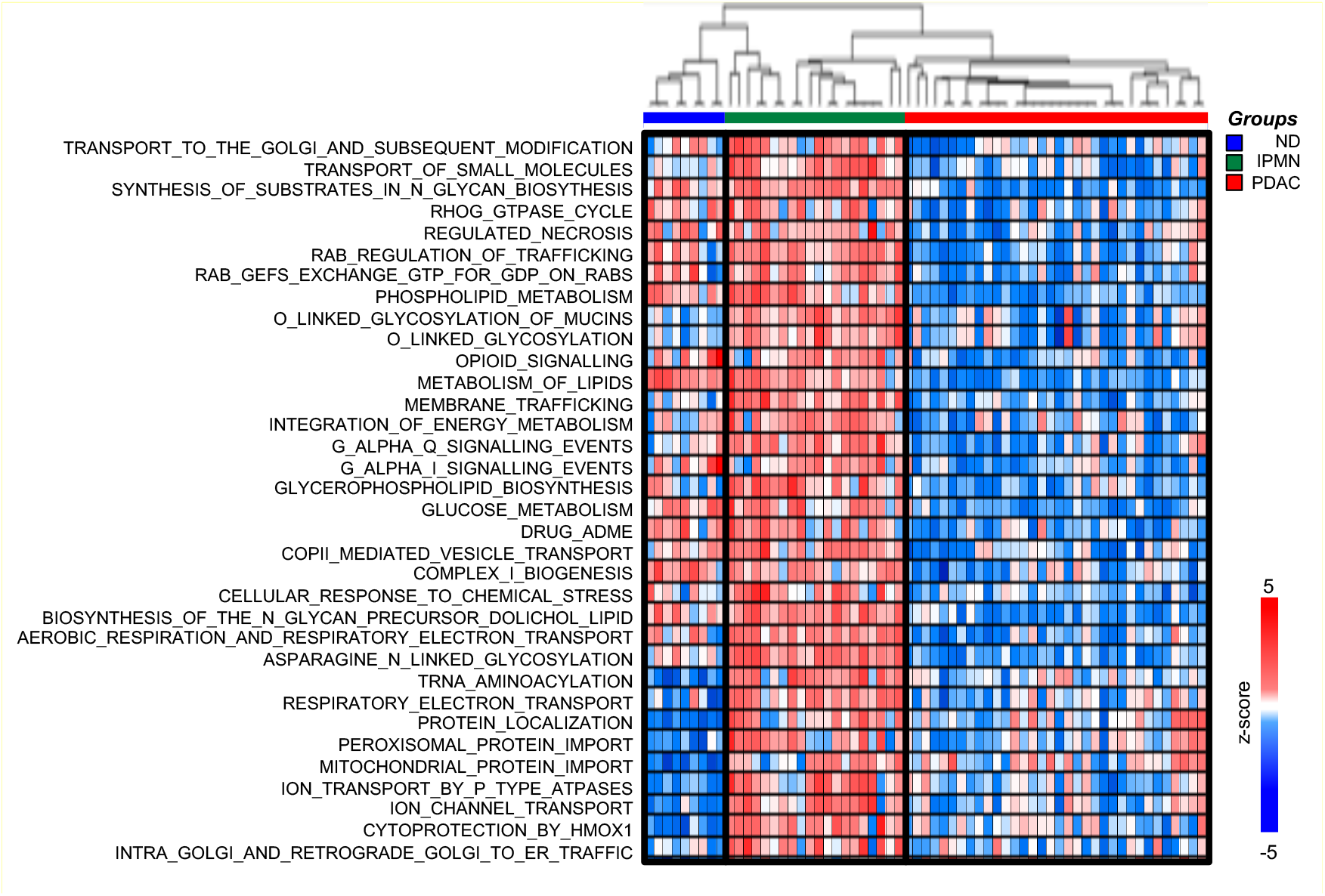
Heatmap illustrating Reactome pathway activation scores from ssGSEA for pathways upregulated in IPMN comparing to normal ducts but downregulated in PDAC. Rows represent individual Reactome pathways, while columns correspond to individual samples grouped by disease condition (ND in blue, IPMN in green, and PDAC in red). The color scale indicates z-scores, with red representing higher activation and blue representing lower activation.

*Table S1. List of proteins identified from pancreas tissue sections subjected to different staining conditions (Hematoxylin, Toluidine Blue O, or unstained). The table provides comparative proteomic profiling across staining methods to evaluate their impact on protein detection in spatial proteomics workflows*.

*Table S2. Protein identification results from laser micro dissected tissue regions captured at different thicknesses (5 μm, 10 μm, and 20 μm) and varying sample sizes representing various cell numbers. The table presents proteomic depth of proteins achieved across conditions, informing the optimization of sample preparation for FFPE-based spatial proteomics*.

*Table S3. Proteins identified from spatially defined tissue regions corresponding to normal duct (ND), intraductal papillary mucinous neoplasm (IPMN), and pancreatic ductal adenocarcinoma (PDAC). This dataset supports comparative analyses of proteomic differences across distinct disease stages using SP-Max workflow*.

*Table S4. Differentially expressed proteins identified via ANOVA (FDR < 0*.*01, S0 = 0*.*1) across ND, IPMN, and PDAC tissue sections. Proteins are categorized into distinct expression clusters, highlighting key molecular transitions during pancreatic cancer progression*.

*Table S5. Biological process enrichment analysis of significantly regulated proteins in IPMN and PDAC. This table presents enriched Gene Ontology (GO) terms associated with upregulated and downregulated protein clusters, shedding light on pathways implicated in pancreatic cancer development*.

*Table S6. Single-sample Gene Set Enrichment Analysis (ssGSEA) results for Reactome pathway activity across ND, IPMN, and PDAC samples. The table quantifies pathway enrichment scores, providing insights into metabolic shifts, immune regulation, and extracellular matrix remodeling during pancreatic tumorigenesis*.

## Data Availability

The mass spectrometry proteomics data have been deposited to the ProteomeXchange Consortium via the PRIDE partner repository with the dataset identifier PXD060378^47^.

Log in to the PRIDE website using the following details:

Project accession: PXD060378

Token: O32CavJ4Uq2e

Alternatively, reviewer can access the dataset by logging in to the PRIDE website using the following account details:

Username

Password: pxTPK6j6dy7h

## Acknowledgments

This work was supported by National Institutes of Health, Pancreatic Cancer Detection Consortium (PCDC, U01CA274514), Clinical Proteomic Tumor Analysis Consortium (CPTAC, U24CA271079), and Early Detection Research Network (EDRN, U2CCA271895).

## Authors Contribution Statement

**J.W**.: Conceptualization, Methodology, Data curation, Writing original draft, visualization **Z.S**.: Methodology **Y.H**.: Data curation **T.H**.: Methodology **D.C**.: Resources **S.K**.: Resources **S.A**.: Resources **B.R**.: Resources **D.C**.: Funding acquisition **K.L**.: methodology, Resources **R.H**.: Resources **H.Z**.: Conceptualization, Supervision, Funding acquisition

## References

(1) Li, B., Zhang, Q., Castaneda, C., Cook, S. Targeted Therapies in Pancreatic Cancer: A New Era of Precision Medicine. Biomedicines 2024, 12 (10). DOI: 10.3390/biomedicines12102175 From NLM PubMed-not-MEDLINE.

(2) Rahib, L., Smith, B. D., Aizenberg, R., Rosenzweig, A. B., Fleshman, J. M., Matrisian, L. M. Projecting cancer incidence and deaths to 2030: the unexpected burden of thyroid, liver, and pancreas cancers in the United States. Cancer Res 2014, 74 (11), 2913–2921. DOI: 10.1158/0008-5472.CAN-14-0155 From NLM Medline.

(3) Quante, A. S., Ming, C., Rottmann, M., Engel, J., Boeck, S., Heinemann, V., Westphalen, C. B., Strauch, K. Projections of cancer incidence and cancer-related deaths in Germany by 2020 and 2030. Cancer Med 2016, 5 (9), 2649–2656. DOI: 10.1002/cam4.767 From NLM Medline.

(4) Prattico, F., Garajova, I. Focus on Pancreatic Cancer Microenvironment. Curr Oncol 2024, 31 (8), 4241–4260. DOI: 10.3390/curroncol31080316 From NLM Medline.

(5) Hegazi, A., Rager, L. E., Watkins, D. E., Su, K. H. Advancing Immunotherapy in Pancreatic Cancer. International journal of molecular sciences 2024, 25 (21). DOI: 10.3390/ijms252111560 From NLM Medline.

(6) Xu, Y., Wang, Y., Hoti, N., Clark, D. J., Chen, S. Y., Zhang, H. The next “sweet” spot for pancreatic ductal adenocarcinoma: Glycoprotein for early detection. Mass Spectrom Rev 2023, 42 (2), 822–843. DOI: 10.1002/mas.21748 From NLM Medline.

(7) Pereira, S. P., Oldfield, L., Ney, A., Hart, P. A., Keane, M. G., Pandol, S. J., Li, D., Greenhalf, W., Jeon, C. Y., Koay, E. J., et al. Early detection of pancreatic cancer. Lancet Gastroenterol Hepatol 2020, 5 (7), 698–710. DOI: 10.1016/S2468-1253(19)30416-9 From NLM Medline.

(8) Wang, Y., Lih, T. M., Lee, J. W., Ohtsuka, T., Hozaka, Y., Mino-Kenudson, M., Adsay, N. V., Luchini, C., Scarpa, A., Maker, A. V., et al. Multi-omic profiling of intraductal papillary neoplasms of the pancreas reveals distinct expression patterns and potential markers of progression. bioRxiv 2024, 2024.2007.2007.602385. DOI: 10.1101/2024.07.07.602385.

(9) Clark, D. J., Zhang, H. Proteomic approaches for characterizing renal cell carcinoma. Clin Proteomics 2020, 17, 28. DOI: 10.1186/s12014-020-09291-w From NLM PubMed-not-MEDLINE.

(10) Cao, L., Huang, C., Cui Zhou, D., Hu, Y., Lih, T. M., Savage, S. R., Krug, K., Clark, D. J., Schnaubelt, M., Chen, L., et al. Proteogenomic characterization of pancreatic ductal adenocarcinoma. Cell 2021, 184 (19), 5031–5052 e5026. DOI: 10.1016/j.cell.2021.08.023 From NLM Medline.

(11) Lih, T. M., Cao, L., Minoo, P., Omenn, G. S., Hruban, R. H., Chan, D. W., Bathe, O. F., Zhang, H. Detection of Pancreatic Ductal Adenocarcinoma-Associated Proteins in Serum. Mol Cell Proteomics 2024, 23 (1), 100687. DOI: 10.1016/j.mcpro.2023.100687 From NLM Medline.

(12) Li, Q. K., Hu, Y., Chen, L., Schnaubelt, M., Cui Zhou, D., Li, Y., Lu, R. J., Thiagarajan, M., Hostetter, G., Newton, C. J., et al. Neoplastic cell enrichment of tumor tissues using coring and laser microdissection for proteomic and genomic analyses of pancreatic ductal adenocarcinoma. Clin Proteomics 2022, 19 (1), 36. DOI: 10.1186/s12014-022-09373-x From NLM PubMed-not-MEDLINE.

(13) Xu, Y., Lih, T. M., De Marzo, A. M., Li, Q. K., Zhang, H. SPOT: spatial proteomics through on-site tissue-protein-labeling. Clin Proteomics 2024, 21 (1), 60. DOI: 10.1186/s12014-024-09505-5 From NLM PubMed-not-MEDLINE.

(14) Woo, J., Schoenfeld, M., Sun, X., Iraguha, T., Zhou, Z., Zhang, Q. Mouse Paneth Cell-Enriched Proteome Enabled by Laser Capture Microdissection. J Proteome Res 2022, 21 (10), 2435–2442. DOI: 10.1021/acs.jproteome.2c00311.

(15) Handa, T., Sasaki, H., Takao, M., Tano, M., Uchida, Y. Proteomics-based investigation of cerebrovascular molecular mechanisms in cerebral amyloid angiopathy by the FFPE-LMD-PCT-SWATH method. Fluids Barriers CNS 2022, 19 (1), 56. DOI: 10.1186/s12987-022-00351-x From NLM Medline.

(16) Mao, Y., Wang, X., Huang, P., Tian, R. Spatial proteomics for understanding the tissue microenvironment. Analyst 2021, 146 (12), 3777–3798. DOI: 10.1039/d1an00472g From NLM Medline.

(17) Bateman, N. W., Conrads, T. P. Recent advances and opportunities in proteomic analyses of tumour heterogeneity. J Pathol 2018, 244 (5), 628–637. DOI: 10.1002/path.5036 From NLM Medline.

(18) Griesser, E., Wyatt, H., Ten Have, S., Stierstorfer, B., Lenter, M., Lamond, A. I. Quantitative Profiling of the Human Substantia Nigra Proteome from Laser-capture Microdissected FFPE Tissue. Mol Cell Proteomics 2020, 19 (5), 839–851. DOI: 10.1074/mcp.RA119.001889 From NLM Medline.

(19) Huang, P., Kong, Q., Gao, W., Chu, B., Li, H., Mao, Y., Cai, Z., Xu, R., Tian, R. Spatial proteome profiling by immunohistochemistry-based laser capture microdissection and data-independent acquisition proteomics. Anal Chim Acta 2020, 1127, 140–148. DOI: 10.1016/j.aca.2020.06.049 From NLM Medline.

(20) Rosenberger, F. A., Thielert, M., Strauss, M. T., Schweizer, L., Ammar, C., Madler, S. C., Metousis, A., Skowronek, P., Wahle, M., Madden, K., et al. Spatial single-cell mass spectrometry defines zonation of the hepatocyte proteome. Nat Methods 2023, 20 (10), 1530–1536. DOI: 10.1038/s41592-023-02007-6 From NLM Medline.

(21) Woo, J., Loycano, M., Amanullah, M., Qian, J., Amend, S. R., Pienta, K. J., Zhang, H. Single-Cell Proteomic Characterization of Drug-Resistant Prostate Cancer Cells Reveals Molecular Signatures Associated with Morphological Changes. Mol Cell Proteomics 2025, 100949. DOI: 10.1016/j.mcpro.2025.100949 From NLM Publisher.

(22) Kitata, R. B., Velickovic, M., Xu, Z., Zhao, R., Scholten, D., Chu, R. K., Orton, D. J., Chrisler, W. B., Zhang, T., Mathews, J. V., et al. Robust collection and processing for label-free single voxel proteomics. Nat Commun 2025, 16 (1), 547. DOI: 10.1038/s41467-024-54643-x From NLM Medline.

(23) Zhou, Y., Zhou, B., Pache, L., Chang, M., Khodabakhshi, A. H., Tanaseichuk, O., Benner, C., Chanda, S. K. Metascape provides a biologist-oriented resource for the analysis of systems-level datasets. Nat Commun 2019, 10 (1), 1523. DOI: 10.1038/s41467-019-09234-6 From NLM Medline.

(24) Subramanian, A., Tamayo, P., Mootha, V. K., Mukherjee, S., Ebert, B. L., Gillette, M. A., Paulovich, A., Pomeroy, S. L., Golub, T. R., Lander, E. S., et al. Gene set enrichment analysis: a knowledge-based approach for interpreting genome-wide expression profiles. Proc Natl Acad Sci U S A 2005, 102 (43), 15545–15550. DOI: 10.1073/pnas.0506580102 From NLM Medline.

(25) Barbie, D. A., Tamayo, P., Boehm, J. S., Kim, S. Y., Moody, S. E., Dunn, I. F., Schinzel, A. C., Sandy, P., Meylan, E., Scholl, C., et al. Systematic RNA interference reveals that oncogenic KRAS-driven cancers require TBK1. Nature 2009, 462 (7269), 108–112. DOI: 10.1038/nature08460 From NLM Medline.

(26) Muraro, M. J., Dharmadhikari, G., Grun, D., Groen, N., Dielen, T., Jansen, E., van Gurp, L., Engelse, M. A., Carlotti, F., de Koning, E. J., et al. A Single-Cell Transcriptome Atlas of the Human Pancreas. Cell Syst 2016, 3 (4), 385–394 e383. DOI: 10.1016/j.cels.2016.09.002 From NLM Medline.

(27) Fu, W., Lin, Y., Bai, M., Yao, J., Huang, C., Gao, L., Mi, N., Ma, H., Tian, L., Yue, P., et al. Beyond ribosomal function: RPS6 deficiency suppresses cholangiocarcinoma cell growth by disrupting alternative splicing. Acta Pharm Sin B 2024, 14 (9), 3931–3948. DOI: 10.1016/j.apsb.2024.06.028 From NLM PubMed-not-MEDLINE.

(28) Tang, J., Chen, L., Chang, Y., Hang, D., Chen, G., Wang, Y., Feng, L., Xu, M. ZBTB7A interferes with the RPL5-P53 feedback loop and reduces endoplasmic reticulum stress-induced apoptosis of pancreatic cancer cells. Mol Carcinog 2024, 63 (9), 1783–1799. DOI: 10.1002/mc.23772 From NLM Medline.

(29) Xu, C., Lu, Z., Hou, G., Zhu, M. Exploring the function and prognostic value of RPLP0, RPLP1 and RPLP2 expression in lung adenocarcinoma. J Mol Histol 2024, 55 (6), 1079–1091. DOI: 10.1007/s10735-024-10251-z From NLM Medline.

(30) Dong, B., Wang, B., Fan, M., Zhang, J., Zhao, Z. Comprehensive analysis to identify PUS7 as a prognostic biomarker from pan-cancer analysis to osteosarcoma validation. Aging (Albany NY) 2024, 16 (10), 9188–9203. DOI: 10.18632/aging.205863 From NLM Medline.

(31) Lu, Y., Ren, X., Zhou, C., Chen, H., Fan, Y., Wang, C. Overexpression of Ribosomal Protein S6 Kinase A4 (RPS6KA4) Predicts a Poor Prognosis in Hepatocellular Carcinoma Patients: A Study Based on TCGA Samples. Comb Chem High Throughput Screen 2022, 25 (13), 2165–2179. DOI: 10.2174/1386207325666220301105850 From NLM Medline.

(32) Javadov, S., Jang, S., Chapa-Dubocq, X. R., Khuchua, Z., Camara, A. K. Mitochondrial respiratory supercomplexes in mammalian cells: structural versus functional role. J Mol Med (Berl) 2021, 99 (1), 57–73. DOI: 10.1007/s00109-020-02004-8 From NLM Medline.

(33) Hollinshead, K. E. R., Parker, S. J., Eapen, V. V., Encarnacion-Rosado, J., Sohn, A., Oncu, T., Cammer, M., Mancias, J. D., Kimmelman, A. C. Respiratory Supercomplexes Promote Mitochondrial Efficiency and Growth in Severely Hypoxic Pancreatic Cancer. Cell Rep 2020, 33 (1), 108231. DOI: 10.1016/j.celrep.2020.108231 From NLM Medline.

(34) Grasso, D., Zampieri, L. X., Capeloa, T., Van de Velde, J. A., Sonveaux, P. Mitochondria in cancer. Cell Stress 2020, 4 (6), 114–146. DOI: 10.15698/cst2020.06.221 From NLM PubMed-not-MEDLINE.

(35) Mates, J. M., Segura, J. A., Alonso, F. J., Marquez, J. Oxidative stress in apoptosis and cancer: an update. Arch Toxicol 2012, 86 (11), 1649–1665. DOI: 10.1007/s00204-012-0906-3 From NLM Medline.

(36) Pisoschi, A. M., Pop, A., Iordache, F., Stanca, L., Predoi, G., Serban, A. I. Oxidative stress mitigation by antioxidants - An overview on their chemistry and influences on health status. Eur J Med Chem 2021, 209, 112891. DOI: 10.1016/j.ejmech.2020.112891 From NLM Medline.

(37) Bellala, R. S., Chittineedi, P., Llaguno, S. N. S., Mosquera, J. A. N., Mohiddin, G. J., Pandrangi, S. L. Down-Regulation of Cysteine-Glutamate Antiporter in ALDH1A1 Expressing Oral and Breast Cancer Stem Cells Induced Oxidative Stress-Triggered Ferroptosis. J Cancer 2024, 15 (19), 6160–6176. DOI: 10.7150/jca.89429 From NLM PubMed-not-MEDLINE.

(38) Atalay Ekiner, S., Gegotek, A., Skrzydlewska, E. The molecular activity of cannabidiol in the regulation of Nrf2 system interacting with NF-kappaB pathway under oxidative stress. Redox Biol 2022, 57, 102489. DOI: 10.1016/j.redox.2022.102489 From NLM Publisher.

(39) Jeon, H., Sterpi, M., Mo, C., Bteich, F. Claudins: from gatekeepers of epithelial integrity to potential targets in hepato-pancreato-biliary cancers. Front Oncol 2024, 14, 1454882. DOI: 10.3389/fonc.2024.1454882 From NLM PubMed-not-MEDLINE.

(40) Kung, H. C., Yu, J. Targeted therapy for pancreatic ductal adenocarcinoma: Mechanisms and clinical study. MedComm (2020) 2023, 4 (2), e216. DOI: 10.1002/mco2.216 From NLM PubMed-not-MEDLINE.

(41) Grant, T. J., Hua, K., Singh, A. Molecular Pathogenesis of Pancreatic Cancer. Prog Mol Biol Transl Sci 2016, 144, 241–275. DOI: 10.1016/bs.pmbts.2016.09.008 From NLM Medline.

(42) van Roey, R., Brabletz, T., Stemmler, M. P., Armstark, I. Deregulation of Transcription Factor Networks Driving Cell Plasticity and Metastasis in Pancreatic Cancer. Front Cell Dev Biol 2021, 9, 753456. DOI: 10.3389/fcell.2021.753456 From NLM PubMed-not-MEDLINE.

(43) Furukawa, T., Kloppel, G., Volkan Adsay, N., Albores-Saavedra, J., Fukushima, N., Horii, A., Hruban, R. H., Kato, Y., Klimstra, D. S., Longnecker, D. S., et al. Classification of types of intraductal papillary-mucinous neoplasm of the pancreas: a consensus study. Virchows Arch 2005, 447 (5), 794–799. DOI: 10.1007/s00428-005-0039-7 From NLM Medline.

(44) Lih, T. M., Jiao, L., Chen, L., Woo, J., Wang, Y., Zhang, H. AUTO-SP: automated sample preparation for analyzing proteins and protein modifications. bioRxiv 2025, 2025.2001.2008.631960. DOI: 10.1101/2025.01.08.631960.

(45) Wang, D., Li, Y., Ge, H., Ghadban, T., Reeh, M., Gungor, C. The Extracellular Matrix: A Key Accomplice of Cancer Stem Cell Migration, Metastasis Formation, and Drug Resistance in PDAC. Cancers (Basel) 2022, 14 (16). DOI: 10.3390/cancers14163998 From NLM PubMed-not-MEDLINE.

(46) Tian, C., Huang, Y., Clauser, K. R., Rickelt, S., Lau, A. N., Carr, S. A., Vander Heiden, M. G., Hynes, R. O. Suppression of pancreatic ductal adenocarcinoma growth and metastasis by fibrillar collagens produced selectively by tumor cells. Nat Commun 2021, 12 (1), 2328. DOI: 10.1038/s41467-021-22490-9 From NLM Medline.

(47) Deutsch, E. W., Bandeira, N., Perez-Riverol, Y., Sharma, V., Carver, J. J., Mendoza, L., Kundu, D. J., Wang, S., Bandla, C., Kamatchinathan, S., et al. The ProteomeXchange consortium at 10 years: 2023 update. Nucleic Acids Res 2023, 51 (D1), D1539–D1548. DOI: 10.1093/nar/gkac1040 From NLM Medline.

